# Context-specificity in causal signaling networks revealed by phosphoprotein profiling

**DOI:** 10.1101/039636

**Authors:** Steven M. Hill, Nicole K. Nesser, Katie Johnson-Camacho, Mara Jeffress, Aimee Johnson, Chris Boniface, Simon E.F. Spencer, Yiling Lu, Laura M. Heiser, Yancey Lawrence, Nupur T. Pande, James E. Korkola, Joe W. Gray, Gordon B. Mills, Sach Mukherjee, Paul T. Spellman

## Abstract

**Summary:** Signaling networks downstream of receptor tyrosine kinases are among the most extensively studied biological networks. However, it remains unclear whether signaling networks depend on biological context. Signaling networks encode causal influences – and not just correlations – between network components. Here, using a causal framework and systematic time-course assays of signaling proteins, we investigate the context-specificity of signaling networks in a cell line system. We focus on a well-defined set of signaling proteins profiled in four breast cancer cell lines under eight stimulus conditions and inhibition of specific kinases. The data, spanning multiple pathways and comprising approximately 70,000 phosphoprotein and 260,000 protein measurements, provide a wealth of testable, context-specific hypotheses, several of which we validate in independent experiments. Furthermore, the data provide a resource for computational methods development, permitting empirical assessment of causal network learning in a complex, mammalian setting.

## Introduction

The complexity of mammalian receptor tyrosine kinase (RTK) signaling continues to pose challenges for the understanding of physiological processes and changes to such processes that are relevant to disease. Networks, comprising nodes and linking directed edges, are widely used to summarize and reason about signaling. Obviously, signaling systems depend on the concentration and localization of their component molecules, so signaling events may be influenced by genetic and epigenetic context. Indeed, in yeast there are striking examples of signaling links that are modulated by context (Good et al., 2009; Zalatan et al., 2012). In disease biology, and cancer in particular, an improved understanding of signaling in specific contexts may have implications for precision medicine by helping to explain variation in therapeutic response and to inform rational therapeutic strategies.

Much work has gone into elucidating genomic heterogeneity, notably intertumoral heterogeneity in cancer, and it has been shown that this heterogeneity is also manifested at the level of differential expression of components of signaling pathways downstream of RTKs (Akbani et al., 2014; Gerlinger and Swanton, 2010; Nickel et al., 2012; Szerlip et al., 2012). However, differences in average protein abundance (as captured in differential expression or gene set analyses) are conceptually distinct from differences in the causal edge structure of signaling networks, with the latter implying a change in the ability of signals to pass between nodes. Causal relationships are also fundamentally distinct from statistical correlations: if there is a causal edge from node *A* to node *B*, then the abundance of *B* may be changed by intervention on *A*. In contrast, if *A* and *B* are merely correlated, but with no causal edge or pathway between them, intervention on *A* will have no effect on *B,* no matter how strong the correlation (for example, if both nodes are correlated due to co-regulation, but with no sequence of mechanistic events linking them).

Canonical signaling pathways and networks (as described for example in textbooks and online resources) summarize evidence from multiple experiments, conducted in different cell types and growth conditions and therefore such networks are not specific to a particular context. Many well-known links in such networks most likely hold in many contexts and so canonical networks remain a valuable source of insights. However, if causal signaling depends on context then using canonical networks alone will neglect context-specific changes, with implications for reasoning, modeling and prediction. Unbiased ‘interactome’ approaches (e.g. Rolland et al., 2014) continue to expand our view of the space of possible signaling interactions. However, due to the complex nature of genetic, epigenetic and environmental influences on signaling, such approaches cannot in general identify signaling events specific to biological context, e.g. specific to a certain cell type under defined conditions.

Here, we studied context-specific signaling using human cancer cell lines. The data span 32 contexts each defined by the combination of (epi)genetics (breast cancer cell lines MCF7, UACC812, BT20 and BT549) and stimuli. In each of the 32 *(cell line, stimulus)* contexts we carried out time-course experiments using kinase inhibitors as interventions. Reverse-phase protein arrays (RPPAs; Tibes et al. 2006) were then used to interrogate signaling downstream of RTKs. We used more than 150 high-quality antibodies targeting mainly total and phosphorylated proteins (see Table S1). The inhibitors applied in each context allowed elucidation of context-specific causal influences between inhibited and downstream phosphoproteins. We also modeled the data using recently developed methods rooted in probabilistic graphical models to reconstruct context-specific networks intended to capture causal interplay between all measured phosphoproteins (and not just interplay related to inhibited nodes). Our results support the notion of context-specificity in causal signaling. In addition, this paper adds to available resources in two ways:

- **Data resource for causal signaling.** Interventional data are essential to go beyond observational statements about protein expression levels towards an understanding of causal signaling heterogeneity. The experimental design used here, spanning all combinations of context, inhibitor and time, allows for a very wide range of analyses, including, but not limited to, analyses of the kind presented here. Large patient datasets are now available in cancer (see, for example, the TCGA Research Network: http://cancergenome.nih.gov), including functional phosphoprotein assays (Akbani et al., 2014), but systematic temporal profiling under interventions is not currently feasible in such settings. The data presented here complement available patient datasets by providing interventional readouts under defined conditions. Furthermore, the data and analyses provide a wealth of testable hypotheses regarding potentially novel and context-specific signaling links.
- **Computational biology benchmarking.** Network reconstruction has long been a core topic in computational biology but performance with respect to learning of causal links has mainly been benchmarked using simulated data that may not adequately reflect the challenges of real data and relevant biology. A previous study established a synthetic network in yeast that was valuable to the computational biology community as it provided a gold-standard network in a biological model (Cantone et al., 2009). However, the number of nodes was small (five) and the synthetic system most likely does not reflect the complexity of mammalian regulation. The design of our experiments allows for systematic testing of causal network learning in a complex mammalian setting and provides a unique resource for development of computational biology methods. The data presented here were used in the recent HPN-DREAM network inference challenge. The challenge focused on causal networks and the data were used to score more than 2000 submitted networks (full details of the challenge are described in Hill et al., 2016).

## Results

### Causal signaling links and context-specificity

In the causal network formulation we use here, an edge or link has the interpretation that inhibition of the parent node can lead to a change in abundance of the child node that is not via any other node in the network. Figure 1A shows an example where a change is observed in a phosphoprotein C under inhibition of an upstream protein A, but there is no edge between phospho-A and phospho-C because the causal effect of phospho-A on phospho-C occurs via phospho-B. However, if phospho-B were unobserved (or not even known to exist) and therefore not included in the network, in the framework we consider here the network would be defined as containing an edge between phospho-A and phospho-C. This would represent an effect of phospho-A on phospho-C that is physically indirect (since it is mediated via the unobserved node) but nevertheless causal (since intervention on A would change C).

**Figure 1.**
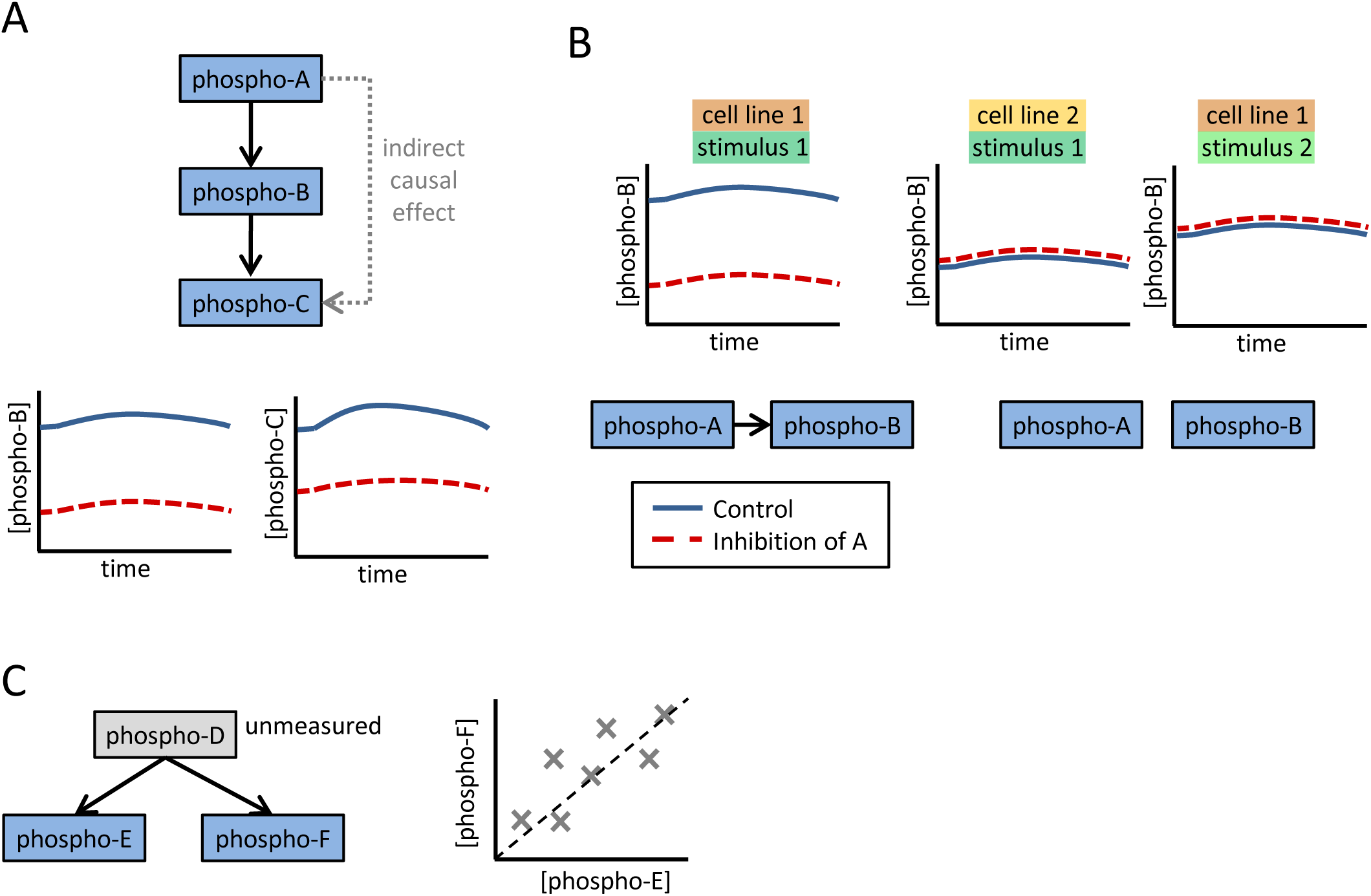
Causal Effects in Signaling (A) An example causal network for a kinase phosphorylation cascade consisting of three phosphoproteins. Phospho-A phosphorylates and activates protein B, which in turn phosphorylates protein C. Due to the influence of phospho-A on protein B, inhibition of A leads to a change in abundance of phospho-B. Although there is no direct physical influence of A on C, such a change is also observed for phospho-C since it is influenced by B, and is therefore a descendant of A in the causal network. (B) Causal edges may be context-specific. In the example shown, there is a causal edge from phospho-A to phospho-B for the *(cell line 1, stimulus 1)* context, but not for *(cell line 2, stimulus 1)* or *(cell line 1, stimulus 2)*. (C) Correlation and causation. Phospho-E and phospho-F are correlated due to regulation by the same protein (phospho-D). However, there is no causal relationship (direct or indirect) between phospho-E and phospho-F and inhibition of protein E would not result in a change in phospho-F.

To illustrate the notion of a context-specific causal edge, consider the situation where there is a causal edge from phospho-A to phospho-B in cell line 1 under stimulus 1, and for this reason inhibition of A leads to a change in the level of phospho-B (Figure 1B). In contrast, in two other contexts, there is no (direct or indirect) causal edge from phospho-A to phospho-B and therefore no change in phospho-B under inhibition of A.

Causation concerns changes under intervention and is fundamentally distinct from correlation. For example, consider the scenario where both phospho-E and phospho-F are regulated by phospho-D (possibly indirectly via unmeasured nodes), but there is no sequence of mechanistic events between phospho-E and phospho-F (Figure 1C). In this scenario, phospho-E and phospho-F may be statistically correlated or dependent (due to co-regulation by phospho-D), but intervention on phospho-E would not be able to induce a change to phospho-F and therefore no causal edge should be placed between E and F.

### Overview of approach

We considered four breast cancer cell lines (MCF7, UACC812, BT20 and BT549) derived from distinct epigenetic states and harboring different genomic aberrations (Barretina et al., 2012; Garnett et al., 2012; Heiser et al., 2012; Neve et al., 2006). Each cell line was serum starved for 24 hours and then, at time t=0min stimulated with one of eight different stimuli (Figure 2A). For each *(cell line, stimulus)* context, we carried out RPPA time-course assays consisting of a total of seven time points spanning four hours, and under five different kinase inhibitors plus DMSO as a control (Figure 2A and Experimental Procedures; here we focused on relatively short-term events – the assays included additional, later time points that were not used in our analyses). To ensure that targets of the kinase inhibitors were effectively blocked, cells were treated with inhibitors for two hours before stimulus. Low concentrations of each inhibitor were used to minimize off-target effects (see Experimental Procedures). Due to the functional significance of phosphorylation, the analyses presented below focus on the 35 phosphoproteins that were measured in all cell lines (see Experimental Procedures and Table S1). Context-specific changes under intervention were summarized as *causal descendancy matrices* (Figure 2B; see below). Machine learning methods were used to integrate the interventional data with known biology to reconstruct context-specific signaling networks (Figure 2C).

**Figure 2.**
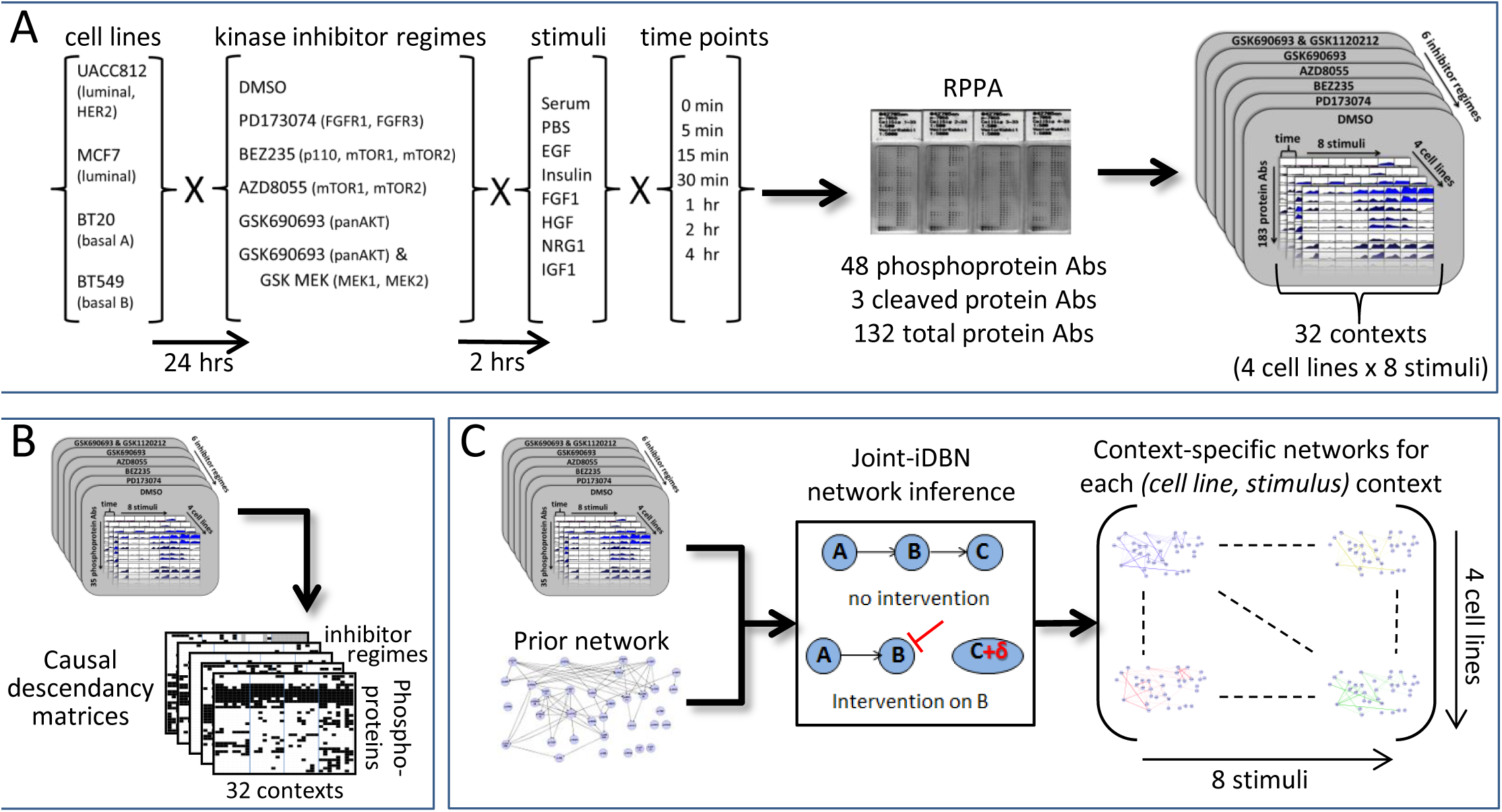
Data-Driven Reconstruction of Context-Specific Causal Signaling Networks.(A) Overview of experimental approach. Reverse-phase protein arrays (RPPA) were used to investigate protein signaling in four human breast cancer cell lines under eight different stimuli. The combinations of cell line and stimulus defined 32 *(cell line, stimulus)* contexts. Prior to stimulus, cell lines were serum starved and treated with kinase inhibitors or DMSO control. RPPA assays were performed for each context at multiple time points post-stimulus, using more than 150 high-quality antibodies to target specific proteins, including approximately 40 phosphoproteins (the precise number of antibodies varies across cell lines; see Experimental Procedures and Table S1). (B) Causal descendancy matrices (CDMs). CDMs summarizing changes under intervention across all contexts were constructed for each intervention. (C) Overview of causal network learning procedure. Interventional time-course data for each context were combined with existing biological knowledge in the form of a prior network to reconstruct context-specific phosphoprotein signaling networks. Networks were learned across all 32 contexts jointly using a variant of dynamic Bayesian networks designed for use with interventional data (see Experimental Procedures).

### Interventional time-course data specific to biological context

Causal signaling links concern changes under intervention (and not merely differential protein expression or changes to statistical correlation between nodes). Comparing time-course data between inhibitor and control (DMSO) experiments allowed us to detect changes to phosphoprotein nodes caused by kinase inhibition (see Experimental Procedures for details). These changes are visualized in a global manner for cell lines UACC812 and MCF7 in Figure 3B, with DMSO time-courses shown in Figure 3A. In Figure 3B, the color coding indicates direction of effect (see examples at bottom right of Figure 3B): green indicates a decrease under intervention relative to control (consistent with positive regulation) and red an increase under intervention (consistent with negative regulation). Corresponding visualizations for BT20 and BT549 are shown in Figure S1.

**Figure 3.**
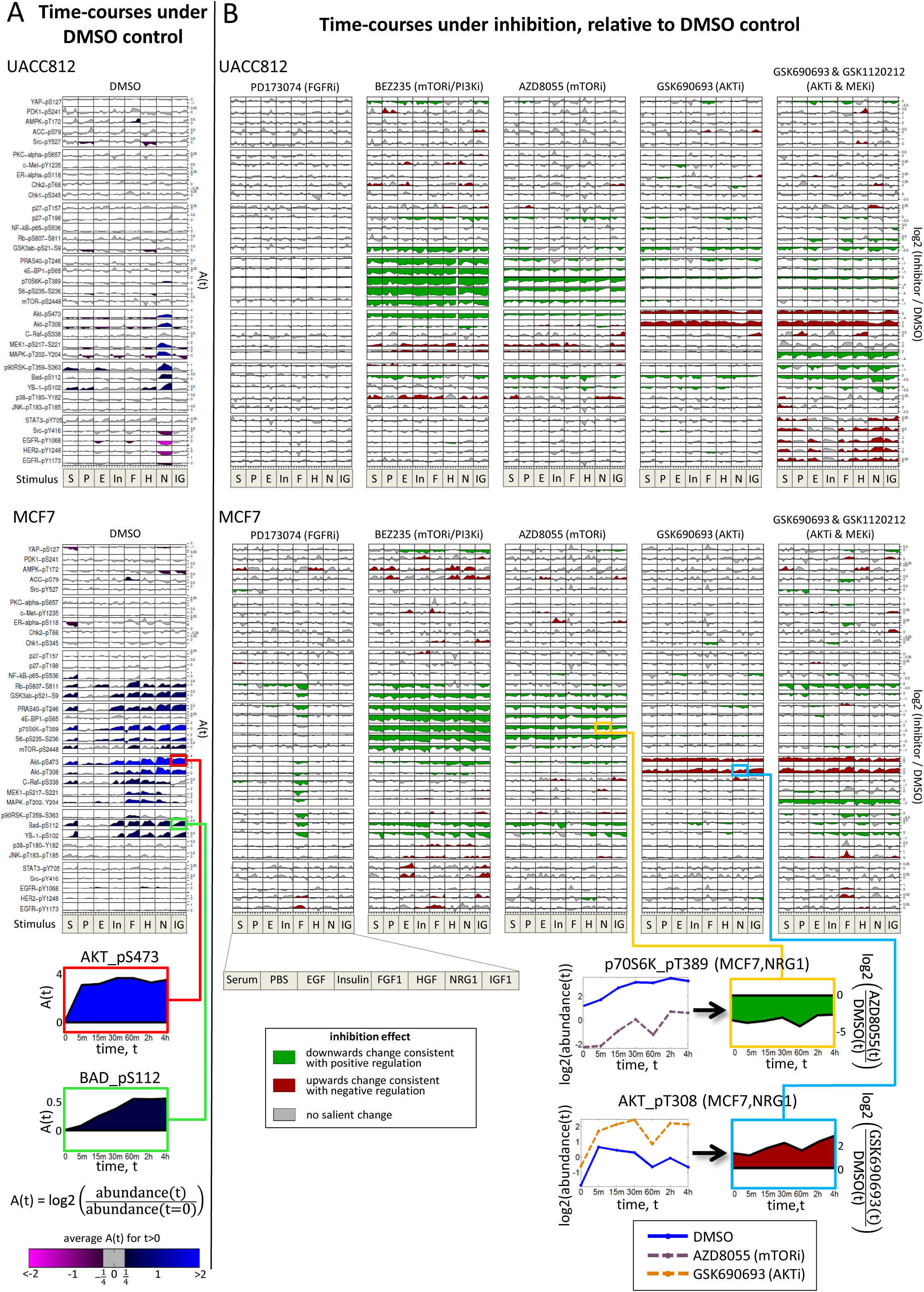
Phosphoprotein Time-Course Data and Context-Specific Changes Under Inhibition for Breast Cancer Cell Lines UACC812 and MCF7. (A) Phosphoprotein time-courses under DMSO control. Rows correspond to 35 phosphoproteins (a subset of the full set of 48; see Experimental Procedures for details) and columns correspond to the eight stimuli. Each time-course shows log_2_ ratios of phosphoprotein abundance relative to abundance at *t = 0*. Shading represents average log_2_ ratio for *t > 0*. (B) Phosphoprotein time-courses under kinase inhibition. Each of the five vertical blocks corresponds to a different inhibition regime. Within each block, rows and columns are as in (A). Each time-course shows log_2_ ratios of phosphoprotein abundance under inhibition relative to abundance under DMSO control. Shading represents direction of changes in abundance due to inhibitor: Green denotes a decrease in abundance (see yellow box example), red denotes an increase (see blue box example) and gray denotes no salient change. See Experimental Procedures for details of statistical analysis. For both (A) and (B), plots were generated using a modified version of the DataRail software (Saez-Rodriguez et al., 2008). Each phosphoprotein is plotted on its own scale and phosphoproteins are ordered by hierarchical clustering of all data. See Figure S1 for corresponding plots for cell lines BT549 and BT20.

Many effects, including many classical ones, are not stimulus-dependent. For example, phospho-p70S6K is reduced relative to control under mTOR inhibition (inhibitor AZD8055; Figure 3B, bottom right), in line with the known casual role of mTOR in regulating phosphorylation of p70S6K. It is important to note that since mTOR signaling is already active in serum starved cells (and not substantially triggered under any stimulus) the reduction in phospho-p70S6K under mTOR inhibition is seen at all time points, including *t*=0min. However, some changes under intervention are specific to individual stimuli. Some of these effects can be readily explained, such as the reduction in abundance of several phosphoproteins in the AKT and MAPK pathways under FGFR inhibition (inhibitor PD173074) for cell line MCF7 stimulated with FGF1. Other stimulus-specific changes are less expected, including the decrease in abundance of phospho-AKT (phosphorylated at threonine 308) in cell line MCF7 under inhibition of mTOR/PI3K (inhibitor BEZ235) that is observed in only four of the stimuli.

### Causal descendancy matrices summarize changes under intervention across multiple contexts

Changes seen under inhibition of mTOR (catalytic inhibitor AZD8055) are summarized in Figure 4A (with phosphoproteins in rows and the 32 contexts in columns). Here, a filled-in box for phosphoprotein *p* in context *c* indicates a salient change under mTOR inhibition (see Experimental Procedures), consistent with a causal effect of mTOR on phosphoprotein *p* in context *c*. This effect could occur via a causal pathway involving other (measured or unmeasured) nodes. In other words, an entry in location (*p,c*) in the matrix indicates that phosphoprotein *p* is a *descendant* of mTOR in the causal signaling network for context *c*; we therefore refer to this matrix as a *causal descendancy matrix* for mTOR. For comparison, an additional column shows proteins that are descendants of mTOR according to a canonical signaling network (Figure 4B; Experimental Procedures). Many of the stimulus-wide classical signaling links mentioned above are also conserved across cell lines. But there are also many examples of causal influences that are both non-canonical and cell line-specific; for example, the abundance of phospho-p38 is elevated in UACC812 cells treated with the mTOR inhibitor AZD8055 under serum stimulation whereas there is no change in phospho-p38 levels in cell line BT549 under the same conditions. Similarly, we obtained causal descendancy matrices for each of the other inhibitors in our study (Figure S2).

**Figure 4.**
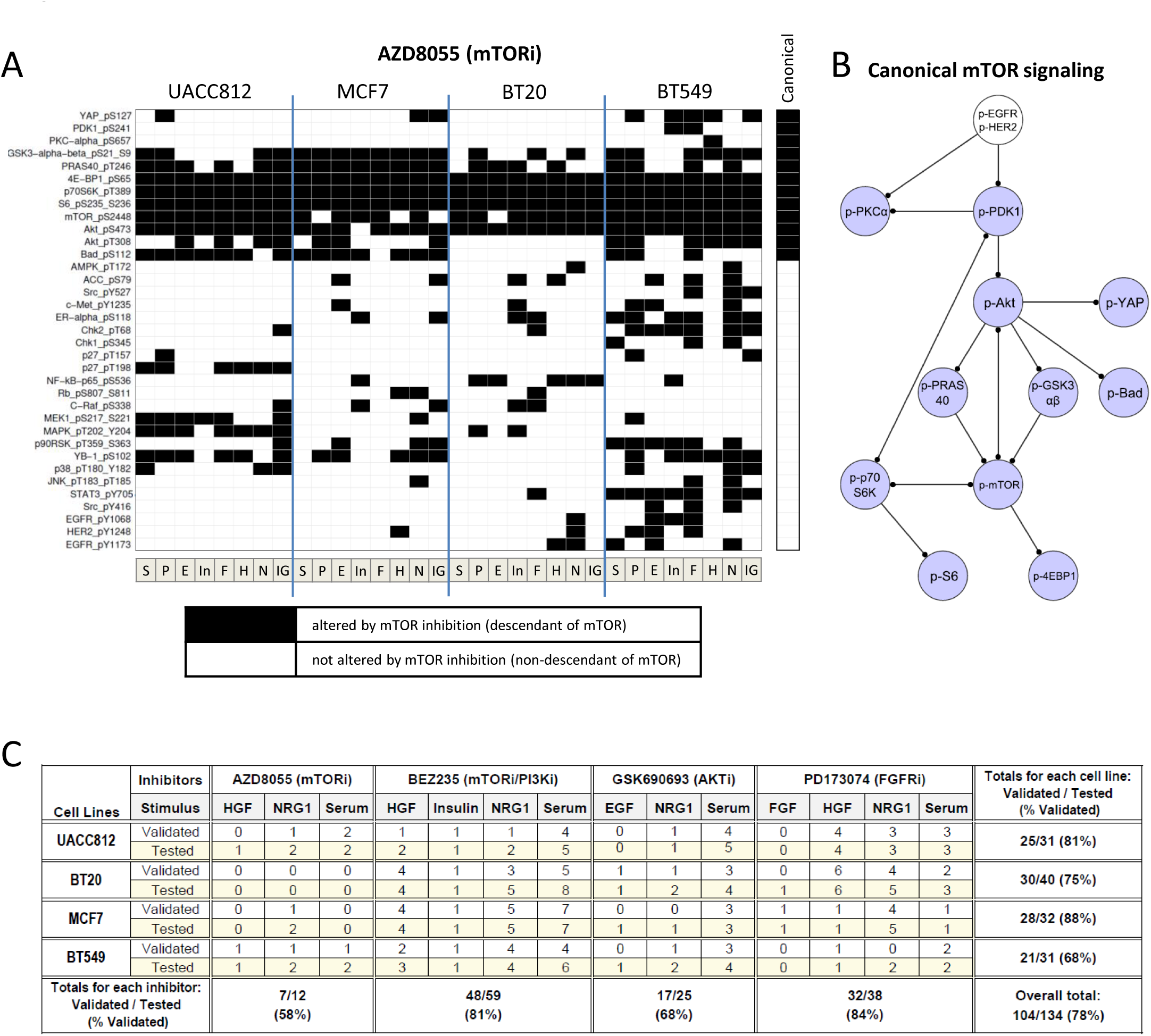
Non-Canonical and Context-Specific Signaling. (A) Causal descendancy matrix showing causal effects observed under mTOR inhibitor AZD8055 in each of the 32 *(cell line, stimulus)* contexts. Rows represent phosphoproteins and columns represent contexts (see Figure 3). Black boxes indicate phosphoproteins that show a salient change under mTOR inhibition in a given context (see Experimental Procedures) and can therefore be regarded as causal descendants of mTOR in the signaling network for that context. The final column on the right indicates phosphoproteins that are descendants of mTOR in the canonical mTOR signaling pathway shown in (B). Phosphoproteins are ordered first by canonical column and then by hierarchical clustering of all data. See Figure S2 for causal descendancy matrices for the other inhibitor regimes. (B) Canonical mTOR signaling pathway. Blue nodes are descendants of mTOR in the network and white nodes are non-descendants. The pathway shown is a subnetwork of the prior network used within the network inference procedure (Figure S3). (C) Summary of western blot validations of causal effects observed in RPPA data. A number of observations from the RPPA data were chosen for validation via western blot analysis. The number of phosphoprotein validations attempted (‘Tested’) and the number of these that successfully validated (‘Validated’) are presented for various *(cell line, stimulus, inhibitor*) combinations. Summary totals are also presented for each cell line, each inhibitor and across all validation experiments. See also Table S2.

We sought to validate some of the observed causal effects by western blot analysis (Experimental Procedures). Observations were selected for validation across both inhibitors and antibodies, and included instances of increase and decrease under inhibition, as well as instances where no effect was observed (Table S2). A summary of the number of observations tested for each cell line and inhibitor regime, and of validation success rate in independent experiments (i.e. new lysates) is shown in Figure 4C.Overall we validated 78% of observations tested (104 out of 134 observations). There were 25 *(antibody, inhibitor)* combinations that for the same stimulus showed differing effects across cell lines in the RPPA data (and which were also tested by western blotting); 17 of these instances of heterogeneity across cell lines validated (68%). The corresponding validation rate for *(antibody, inhibitor)* combinations that for the same cell line showed differing effects across stimuli was only 3 out of 13 (23%). The inability to confirm some RPPA data observations with western blotting could represent biological variability, differential sensitivity between RPPA and western blotting, use of different antibodies or other technical issues.

### Machine learning of signaling networks

We used a machine learning approach to learn context-specific causal networks over all measured phosphoprotein nodes (including those not intervened upon). Specifically we used a variant of dynamic Bayesian networks (see Experimental Procedures and references therein for details) that modeled the interventional time-course data to learn networks simultaneously across all 32 contexts, whilst taking account of known biology (encoded as a prior network, Figure S3).

Figure 5 summarizes networks across all 32 *(cell line, stimulus)* contexts. Here, we see that while many edges, including several classical signaling relationships, are near universal, others are cell line-specific, mirroring, via a global analysis, the inhibition data reported above (Figure 4A). The networks contained edges included in the prior network as well as many edges that were not. Across the 32 contexts, networks contained an average of 49 edges (at a threshold of 0.2 applied to the edge probabilities that are the output of the learning procedure) and, on average, 40% of edges in each network were not in the prior network (Table S3). We discuss potentially novel edges that were not in the prior (and their validation) below. As discussed in Hill et al. (2016), the challenging nature of causal network learning means that empirical performance assessment is important. We used the train-and-test procedure described in Hill et al. (2016) to systematically assess causal network learning (see Experimental Procedures). We found that the models were able to achieve significant agreement with unseen test interventional data in most of the contexts (Figure S4). However, we note that such empirical assessment is an open area in causal inference and the assessment procedure used here is subject to a number of caveats (see Discussion).

**Figure 5.**
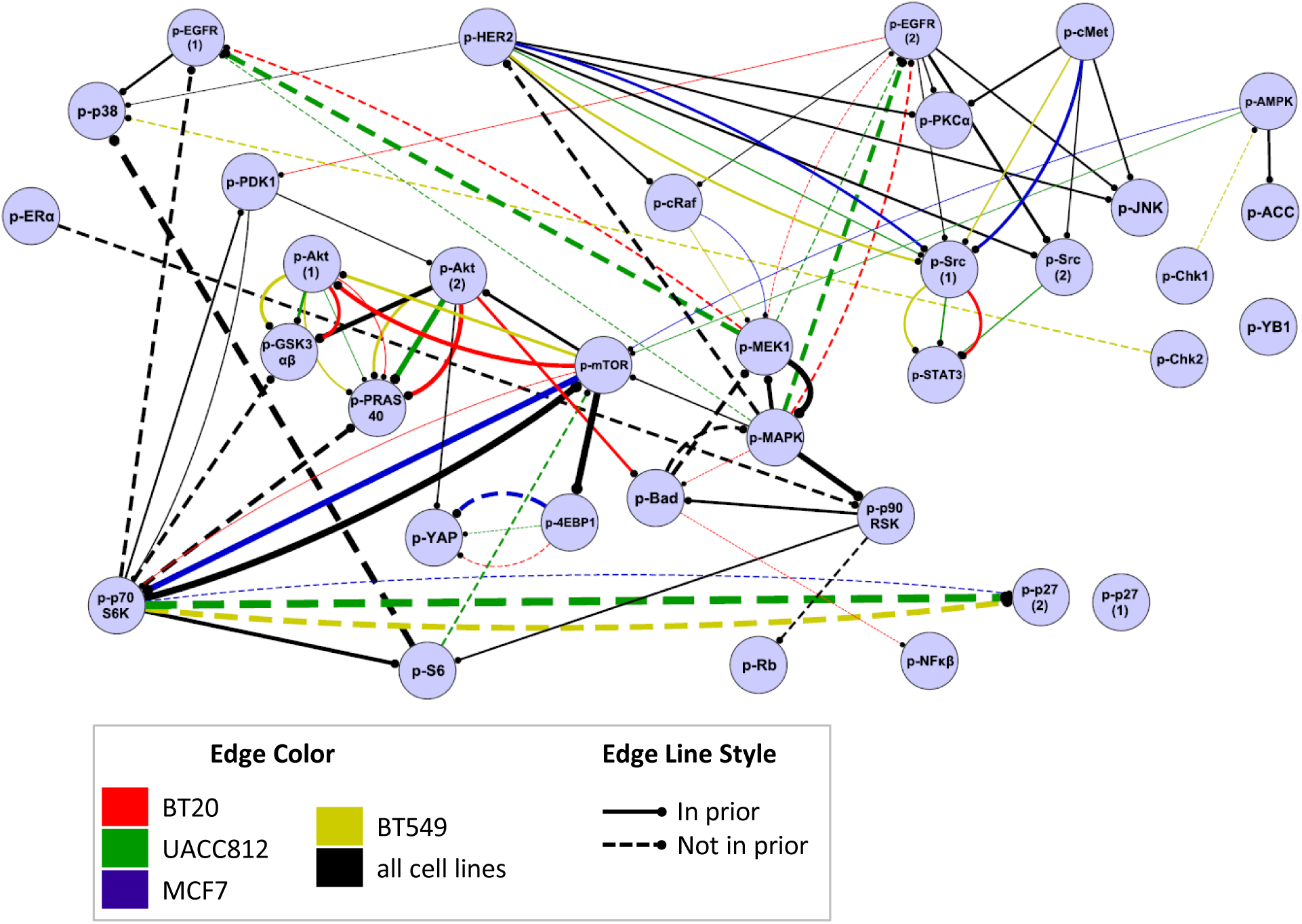
Context-Specific Signaling Networks Reconstructed Using a Machine Learning Approach. Data for 35 phosphoproteins were analyzed using a machine learning approach based on interventional dynamic Bayesian networks, integrating also known biology in the form of a prior network. This gave a set of scores (edge probabilities) for each possible edge in each *(cell line, stimulus)* context (see Experimental Procedures). For each cell line, a summary network was obtained by averaging edge probability scores across the eight stimulus-specific networks for that cell line. Edge color denotes cell line. Only edges with average probabilities greater than 0.2 are shown. A black edge indicates an edge that appears (i.e. is above the 0.2 threshold) in all four cell lines. Edge thickness is proportional to the average edge probability (average taken across all 32 contexts for black edges). Solid/dashed edges were present/not present in the prior network respectively. Edges are directed with the child node indicated by a circle. Edge signs are not reported; the modeling approach does not distinguish between excitatory and inhibitory causal effects. Network visualized using Cytoscape (Shannon et al., 2003). See also Figure S3, Table S3 and Table S4.

### Validation of context-specific signaling hypotheses

In the global networks we identified 235 edges that were not in the prior network but that had edge probability scores above a threshold of 0.2. These potentially novel edges shared 35 parent proteins, 4 of which were inhibited in the original dataset. Five edges with parent nodes not among those inhibited in the original RPPA data were selected for validation by western blot. Edge selection was done on the basis of biological interest and availability of sufficiently specific inhibitors for the parent nodes (Figure 6). We note that our computational approach predicts presence/absence of each edge and its direction, but not sign (activating or inhibiting).

**Figure 6.**
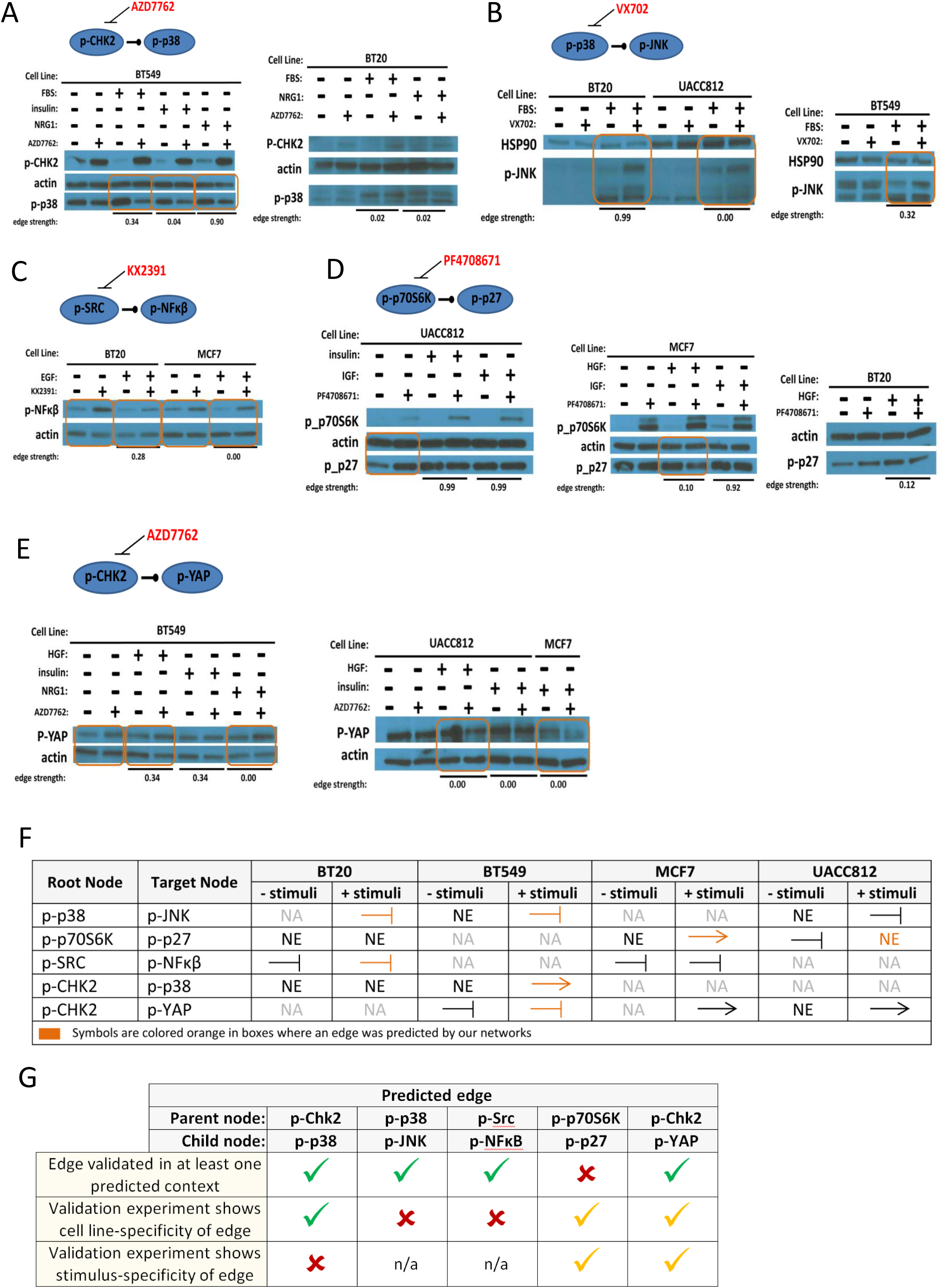
Validation of Novel Network Edges Western blot analysis to validate selected context-specific network edges that were not in the prior network. Edges tested were: (A) phospho-Chk2 to phospho-p38; (B) phospho-p38 to phospho-JNK; (C) phospho-Src to phospho-NFκB; (D) phospho-p70S6K to phospho-p27; and (E) phospho-Chk2 to phospho-YAP. Orange boxed areas indicate observed changes in abundance of the predicted child node under inhibition of the parent node in a single *(cell line, stimulus)* context. Edge probabilities output by the network learning procedure are shown for each context tested (‘edge strength’). (F) A summary of the validation experiments. ‘NA’ denotes ‘not applicable’ – the experiment was not run. ‘NE’ denotes ‘no edge’ – there was no change in child node abundance upon inhibition of the parent node. An arrow indicates results consistent with an activating parent node. A stunted line represents results consistent with an inhibitory edge. Symbols are colored orange to indicate that an edge was predicted for the corresponding cell line under one of the stimuli tested. (G) Summary of agreement and disagreement between predicted edges and validation experiments. First row indicates whether validation experiments showed evidence for the edge in a *(cell line, stimulus)* context in which it was predicted. Second and third rows concern the cell line‐ and stimulus-specificity of each edge respectively: a green tick denotes specificity in (partial) agreement with predictions from inferred networks; an orange tick denotes specificity, but not in agreement with predictions in terms of the precise contexts in which effects were seen; a red cross indicates that specificity was not observed in the validation experiments, despite being predicted by the networks.

For each of the five edges we tested contexts in which the edge was predicted as well as those in which the edge was not predicted. We inhibited the parent node and observed whether this altered abundance of the predicted child node. We found evidence supporting each of the five predicted causal edges, but with often complex context-dependence. These results – and their agreement and disagreement with context-specific predictions from network modeling – are summarized in Figure 6F,G.

A novel edge from Chk2_pT68 to p38_pT180/Y182 (for phosphoproteins we give the protein name before an underscore which is followed by the phosphorylation site(s)) was predicted only in cell line BT549 (Figure 5). When Chk2 is inhibited with AZD7762 in BT549 cells, phospho-p38 decreases under FBS and NRG1 (where the edge was predicted) as well as under insulin (where the edge was not predicted; Figure 6A). In contrast, there is no change in phospho-p38 in BT20 cells under AZD7762 treatment, consistent with the absence of the edge in the BT20 networks. Here we see that the edge validates in a cell line-specific but not stimulus-specific manner. However, it is important to note that AZD7762 inhibits Chk1 and Chk2 with equal potency and also demonstrates activity, albeit lower, against other kinases. Whether this explains why AZD7762 demonstrated activity in insulin treated cells, which was not predicted by the model, remains to be established.

The networks predicted an edge from p38_pT180/Y182 to JNK_pT183/T185 in BT549 and BT20 cells (under stimulus with FBS) and in BT549 cells (under HGF). Upon inhibition of p38 with VX702 in BT20 and BT549 cells stimulated with FBS, we observe an increase in phospho-JNK (Figure 6B). We also observe a modest increase in phospho-JNK in UACC812 cells, although the phospho-p38 to phospho-JNK edge was not predicted in UACC812.

An edge from Src_p416 to NFκβ-p65_pS536 was predicted only in BT20 cells stimulated with EGF. Upon inhibition of Src with KX2391 both before and after stimulation with EGF, an increase in the abundance of phospho-NFκβ was observed in BT20 cells, consistent with the presence of a causal link (Figure 6C). The connection between phospho-Src and phospho-NFκβ was also observed in MCF7, where the edge was not predicted.

An edge from p70S6K_pT389 to p27_pT198 was predicted in all of the UACC812 and BT549 networks. The edge was also predicted in MCF7 networks for PBS, insulin, FGF, NRG1, and IGF1 and in the BT20 NRG1 network. When p70S6K was inhibited in UACC812 cells with PF4708671, a change in phospho-p27 was observed only at the zero time point before stimulus was added (Figure 6D). In MCF7 cells stimulated with HGF, phospho-p27 decreased in abundance under p70S6K inhibition; however, the edge was not predicted in this context. When PF4708671-treated MCF7 cells were stimulated with IGF, a context in which the edge was predicted with high probability, no change in phospho-p27 was observed. Similarly, there was no change in phospho-p27 in BT20 cells that had been treated with PF4708671 and stimulated with HGF.

In BT549 an edge was predicted from Chk2_pT68 to YAP_pS127 under HGF and insulin. BT549 cells treated with the Chk2 inhibitor AZD7762 exhibit an increase in phospho-YAP (Figure 6E). This edge was not predicted in any other cell line tested. However, in both UACC812 and MCF7 cells treated with AZD7762, a decrease in the abundance of phospho-YAP is observed. Active Chk2 appears to decrease phospho-YAP in BT549 cells (where the edge was predicted) and increase phospho-YAP in UACC812 and MCF7 cells (where the edge was not predicted). These results are consistent with the existence of a causal influence of phospho-Chk2 on phospho-YAP in all of these cell lines, and not just in BT549 as predicted.

Thus, we see evidence for each of the five novel edges, demonstrating the utility of our approach in generating testable hypotheses. Furthermore, the validation data support the notion of differences in causal links between contexts (Figure 6F,G). However, although we see effects in several contexts in which they were predicted, effects are also seen in contexts in which they were not predicted, suggesting that predictions are not accurate in terms of precisely which contexts the effect is specific to.

## Discussion

The data and analyses presented here support the view that causal signaling networks can depend on biological context. We focused on breast cancer cell lines. These represent contexts that are genetically perturbed but with a shared tissue-of-origin. The causal signaling heterogeneity that we observed suggests that substantial differences could exist between, for example, samples from different tissue types or divergent environmental conditions. This strongly argues for a need to refine existing models of signaling for specific contexts, not least in disease biology.

Given the range of potentially relevant contexts – spanning combinations of multiple factors, including genetic, epigenetic and environmental – we do not believe that characterization of causal signaling across multiple contexts can feasibly be done using classical approaches in a protein-by-protein manner. Rather, it will require high-throughput data acquisition and computational analysis. Such a program of research requires an appropriate conceptual framework, rich enough to capture regulatory relationships, but still tractable enough for large-scale investigation. Furthermore, for practical application, such an approach will also need to be sufficiently robust to missing or unknown variables. Causal models may provide an appropriate framework because, unlike purely correlational models, they allow for reasoning about change under intervention and are, to a certain extent, robust to missing variables. In particular, causal descendancy matrices (Figures 4A and S2) are robust to missing variables in the sense that addition of a protein (row) to the matrix would not change the existing entries (since the claim that node *A* has a causal influence on node *B* is consistent with missing intermediate nodes). A systematic program of investigation into context-specific signaling will be important for cancer biology, but perhaps even more so for the many other disease areas in which signaling is influential, but that have to date been less well represented than cancer in signaling studies.

Our results extend the well-established notion of genomic intertumoral heterogeneity in cancer to the level of signaling phenotype. We found that cell line-specific findings were more reliable than stimulus-specific findings. This may be due to the magnitude of epigenetic and genetic differences between cell lines being more marked than differences between stimuli, all of which activated closely related cell surface receptors.

We attempted to validate five predicted edges that were not present in the prior network. All five edges showed causal effects in validation experiments, but only one (phospho-Chk2 to phospho-p38; Figure 6A) in a manner consistent with context-specific predictions. In common with most protein profiling studies, including both low‐ and high-throughput techniques, our experiments were based on bulk assays and can therefore only elucidate signaling heterogeneity at the level of cell populations; we did not consider cell-to-cell heterogeneity, tumor stromal interactions, nor the spatial heterogeneity of tumors that plays an important role *in vivo* (Bedard et al., 2013; González-García et al., 2002). However, our data have implications for inter‐ and intra-tumoral heterogeneity because they suggest the possibility that *in vivo* causal signaling networks – and in turn the cell fates and disease progression events that they influence – may depend on the local micro-environment. Further work will be needed to elucidate such dependence and to draw out its implications.

In the future, signaling models may start to play a role in clinical informatics, for example by helping to inform rational assignment of targeted therapies. An implication of the context-specificity we report is that such analyses may require models that are learned, or at least modified, for individual samples (or subsets of samples). While causal models are in some ways simpler than fully dynamical ones, causal inference remains fundamentally challenging and is very much an open area of research. For this reason, alongside advances in relevant assays, a personalized, network-based approach will require suitable empirical diagnostics, a view that echoes long-standing empirical critiques of causal theory (e.g. Freedman & Humphreys 1999). Hill et al. (2016) used the data presented here to score, in an automated manner, over 2000 networks (∼70 methods each applied to infer 32 context-specific networks) submitted to the HPN-DREAM network inference challenge and we used this assessment procedure here. Such assessment procedures could in the future allow for automated quality control, for example rejecting networks that are not sufficiently consistent with unseen interventional readouts (e.g. we did not obtain statistically significant performance under any test inhibitor for the *(BT549, EGF)* context; see Figure S4). However, as discussed in Hill et al. (2016) the assessment approach remains limited in several ways and this argues for caution in interpreting the relatively good performance with respect to the assessment procedure reported here. Of particular relevance to context-specificity, we note that the assessment procedure focuses on global agreement with held-out interventional data and not specifically on identification of differences between contexts. Indeed, our validation experiments showed that although all novel edges that were tested validated in one or more contexts, network predictions were not accurate with respect to the precise context(s) in which changes were seen.

Recently, Carvunis and Ideker (2014) proposed a view of cellular function involving hierarchies of elements and processes and not just networks. Building detailed dynamical or biophysical models over hierarchies spanning multiple time and spatial scales may prove infeasible. A more tractable approach may be to extend coarser causal models of the kind used here in a hierarchical direction, for example by allowing causal links to cross scales and subsystems. Thus, the approach we pursued – of causal models based on context-specific perturbation data – could in the future be used to populate models over cellular hierarchies.

## Experimental Procedures

### Preparation of RPPA samples

Breast epithelial cells in log-phase of growth were harvested, diluted in the appropriate media (RPMI or DMEM) containing 10% fetal bovine serum, and then seeded into 6 well plates at an optimized cell density. BT20 cells were plated at 230,000 cells/ well; BT549 cells were plated at 175,000 cells/ well; MCF7 cells were plated at 215,000 cells/ well; and UACC812 cells were plated at 510,000 cells/well. After 24 hour growth at 37C and 5% CO2 in complete medium, cells were synchronized by incubating with serum-free medium for an additional 24 hours. The medium was then exchanged with fresh serum-free medium containing either: 15nM AZD8055, 50nM GSK690693, 50nM BEZ235, 150nM PD173074, 10nM GSK1120212 in combination with 50nM GSK690693, or vehicle alone (0.05% DMSO) and incubated for two hours prior to stimulation. Cells were then either harvested (0 time point) or stimulated by addition of 200uL per well of 10X stimulus (either PBS, Fetal Bovine Serum, 100ng/mL EGF, 200ng/mL IGF1, 100nM insulin, 200ng/mL FGF1, 1ug/mL NRG1, or 500 ng/mL HGF) for 0, 5, 15, 30 or 60 minutes, or 2, 4, 12, 24, 48 or 72 hours prior to protein harvest. Cells were washed twice with PBS and lysed by adding 50uL of lysis buffer (1% Triton X-100, 50mM HEPES, pH 7.4, 150mM NaCl, 1.5mM MgCl2, 1mM EGTA, 100mM NaF, 10mM Na pyrophosphate, 1mM Na3VO4, 10% glycerol, containing freshly added protease and phosphatase inhibitors from Roche Applied Science 04693116001 and 04906845001, respectively). Lysates were collected by scraping after 20 minutes incubation on ice. Lysates were spun at 4°C in a tabletop centrifuge at 15,000 RPM for 10 minutes and soluble proteins contained in the supernatant were collected. Protein concentration was determined by the Pierce BCA Protein Assay according to manufacturer’s protocol. Protein was then diluted to 1 mg/mL and 30uL of the diluted lysate was mixed with 10uL 4X SDS sample buffer (40% Glycerol, 8% SDS, 0.25M Tris-HCL, pH 6.8 and 10% v/v 2-mercaptoethanol added fresh) and boiled for 5 minutes prior to freezing and shipment to MD Anderson Cancer Center Functional Proteomics Core Facility (Houston, Texas) for Reverse Phase Protein Array (RPPA) analysis (Tibes et al., 2006).

RPPA methodology has been described previously (see e.g. Akbani et al., 2014) and is also outlined in Extended Experimental Procedures. Antibodies used in the assay go through a validation process as previously described (Hennessy et al., 2010) to assess specificity, quantification and dynamic range. The validation status of each antibody can be found in Table S1.

### Western Blot Analysis

Cells were grown as described above. Additional inhibitors were used to generate lysates for novel edge validations following the protocol laid out above. The inhibitors, all used at 1μM, were AZD7762, KX2-391, PF4708671, and VX-702 (see Figure 6 for targets). Lysates were harvested 15 minutes after stimulation and protein concentrations quantified as described above. Denatured lysates were run on 4–12% Bis-Tris gradient gels (Invitrogen). Gels were transferred to immobilin-FL PVDF membranes (Millipore) before being immunoblotted with indicated antibodies (Table S5).

### Data quality control and pre-processing

Several further quality control and pre-processing steps were performed prior to analysis of the RPPA time-course data. These steps are briefly summarized below, with some additional details provided in Extended Experimental Procedures.

Several samples were excluded from analysis because they did not pass quality control (Table S6). These samples were identified using protein loading correction factors, variance across antibodies, and signal-to-noise ratio analysis (see Extended Experimental Procedures). In addition, data for the combination of inhibitors GSK690693 & GSK1120212 (AKTi & MEKi) for cell lines BT549 (all stimuli) and BT20 (PBS and NRG1 stimuli only) were excluded since none of the expected effects of MEKi were observed in these samples.

A batch normalization procedure was performed for cell line UACC812 due to samples for this cell line being split across two batches. Normalization was performed using samples that were common to both batches and is fully described in Extended Experimental Procedures. Data were log transformed and replicates (only present for *t*=0 samples and some DMSO samples) were averaged. Prior to input into our network inference pipeline, imputation was performed for missing data by linear interpolation of adjacent time points.

To facilitate comparisons between cell lines, the analyses presented here focused on the set of phosphoprotein antibodies common to all four lines. This set contained two highly correlated pairs of antibodies (*r* > 0.9 for all cell lines), consisting of phosphoforms of the same protein: GSK3αβ_pS21_pS9, GSK3_pS9 and S6_pS235_S236, S6_pS240_S244. Only one antibody out of each pair was included in analyses, resulting in a final set of 35 phosphoprotein antibodies. A full list of antibodies can be found in Table S1.

### Identification of changes under kinase inhibition

We used a procedure centered on paired *t*-tests to determine which phosphoproteins show a salient change in abundance under each kinase inhibitor. Details are described in Hill et al. (2016), but also outlined below for completeness.

For each phosphoprotein, inhibitor regime and *(cell line, stimulus)* context, a paired *t*-test was used to assess whether mean phosphoprotein abundance under DMSO control is significantly different to mean abundance under the inhibitor regime (mean values calculated over seven time points). Some phosphoproteins show a clear response to the stimulus under DMSO control, with abundance increasing and then decreasing over time (a ‘peak’ shape), while others show a less clear response due to signal already being present prior to stimulus. For phosphoproteins falling into the former category (according to a heuristic), paired *t*-tests were repeated, but this time restricted to intermediate time points within the peak. This focuses on the portion of the time-course where an inhibition effect, if present, should be seen. The *p*-value from the repeated test was retained if smaller than the original *p*-value. For each *(cell line, stimulus)* context and inhibitor regime, the resulting set of *p*-values (one *p*-value for each phosphoprotein) were corrected for multiple testing using the adaptive linear step-up procedure for controlling the FDR (Benjamini et al., 2006).

For each *(cell line, stimulus)* context, a phosphoprotein was deemed to show a salient change under a given inhibitor regime if two conditions were satisfied. First, the corresponding FDR value had to be less than 5% and, second, the effect size (log_2_ ratio between DMSO control and inhibitor conditions) had to be sufficiently large relative to replicate variation. The latter condition is an additional filter to remove small effects. The phosphoproteins satisfying these criteria are depicted in Figures 3B, 4A, S1 and S2. We note that the overall procedure is heuristic and that the FDR values should not be interpreted formally.

A phosphoprotein *p* showing a salient change under an inhibitor is consistent with a node targeted by the inhibitor having a causal effect on the phosphoprotein. Since this effect can be direct or indirect, phosphoprotein *p* can be regarded as a descendant of the inhibitor target node in the underlying signaling network. That is, there exists a directed path starting from the node targeted by the inhibitor and ending at phosphoprotein *p*.

### Network learning

Networks were learned for each of the 32 *(cell line, stimulus)* contexts using dynamic Bayesian networks (DBNs), a type of probabilistic graphical model for time-course data (see e.g. Hill et al., 2012; Husmeier, 2003; Murphy, 2002). Specifically we used a variant called *interventional DBNs* or iDBNs (Spencer et al., 2015), that uses ideas from causal inference (Pearl, 2009; Spirtes et al., 2000) to model interventions and thereby improve ability to infer causal relationships; model specification followed Spencer et al. (2015). Although interested in learning context-specific networks, we expect a good proportion of agreement between contexts. Therefore, rather than learn networks for each context separately, we used a joint approach to learn all networks together (Oates et al., 2014). A prior network was used (Figure S3); this was curated manually with input from literature (Weinberg, 2013) and online resources. The extent to which context-specific networks are encouraged to agree with each other and with the prior network is controlled by two parameters, λ and η respectively, as described in detail in Oates et al. (2014). These parameters were set (to λ = 3 and η = 15) by considering a grid of possible values and selecting an option that provides a reasonable, but conservative amount of agreement, allowing for discovery of context-specific edges that are not in the canonical prior network. The network learning approach resulted in a score (edge probability) for each possible edge in each context-specific network. The network estimates were robust to moderate data deletion and precise specification of the biological prior network and its strength (Figure S5). Furthermore, the analyses were computationally efficient, requiring approximately 30 minutes to learn all 32 context-specific networks using serial computation on a standard personal computer (Intel i7-2640M 2.80GHz processor, 8GB RAM).

### Assessing performance of causal network learning

The ability of our network learning approach to estimate context-specific causal networks was systematically assessed using a train and test scheme proposed by Hill et al. (2016) in the context of the HPN-DREAM network inference challenge associated with the RPPA data presented here. In brief, networks were learned on a training dataset consisting of a subset of the inhibitors, and the causal validity of the networks was then assessed using held-out data from an unseen, test inhibitor. In contrast to Hill et al. (2016) where, due to factors specific to the challenge setting, a single iteration of train and test was used, we iterated over available inhibitors. Full details of the assessment approach can be found in Hill et al. (2016), but for completeness we also provide an outline in Extended Experimental Procedures.

